# Silversword and lobeliad reintroduction linked to landscape restoration on Mauna Loa and Kīlauea, and its implications for plant adaptive radiation in Hawaiʻi

**DOI:** 10.1101/095216

**Authors:** Robert H. Robichaux, Patrice Y. Moriyasu, Jaime H. Enoka, Sierra McDaniel, Rhonda K. Loh, Kealiʻi F. Bio, Ane Bakutis, J. Timothy Tunison, Steven T. Bergfeld, J. Lyman Perry, Frederick R. Warshauer, Mark Wasser, T. Colleen Cole, Nicholas R. Agorastos, Ian W. Cole, J. Kualiʻi Camara, Tanya Rubenstein, A. Nāmaka Whitehead, Joshua R. VanDeMark, Reid Loo, Marie M. Bruegmann

## Abstract

The endemic Hawaiian silversword and lobeliad lineages, which are two of the world’s premier examples of plant adaptive radiation, exemplify the severity of the threats confronting the Hawaiian flora, especially the threats posed by alien species. We have implemented collaborative reintroduction efforts with the endangered Kaʻū silversword (*Argyroxiphium kauense*) and Pele lobeliad (*Clermontiapeleana)* in Hawaiʻi Volcanoes National Park. The efforts with the Kaʻū silversword have involved rediscovery, helicopter assisted rescue of diminutive remnant founders, managed breeding, outplanting at two sites in the Park of more than 21,000 seedlings deriving from 169 founders, and facilitated achene dispersal following flowering. The efforts with the Pele lobeliad have involved rediscovery, air-layering of remnant founders while suspended on climbing ropes, managed breeding, and outplanting at two sites in the Park of more than 1,000 seedlings (to date) deriving from six of the seven known founders. We have linked the reintroduction efforts to landscape restoration at large scales in the Park and in adjacent State and private lands, thereby increasing the opportunities for substantial population growth and expansion of the Kaʻū silversword and Pele lobeliad in the future. Additionally, we have extended the reintroduction efforts, including the link to landscape restoration, to encompass all other endangered silversword and lobeliad taxa occurring historically on the eastern slopes of Mauna Loa or on Kīlauea. In so doing, we seek to restore the possibility of adaptive radiation of the silversword and lobeliad lineages going forward, especially on the youngest and most geologically active, and thus perhaps most evolutionarily dynamic, part of the Hawaiian archipelago.

**Dedication:** This paper celebrates the centennial of Hawaiʻi Volcanoes National Park, which was founded in August 1916.

## Introduction

Oceanic archipelagos well-illustrate two key aspects of life’s diversity and conservation: (1) adaptive radiation, which generates great diversity, and (2) human mediated invasion by alien species, which imperils diversity. Adaptive radiation is one of the more spectacular processes in organismal evolution, with single colonizing species giving rise to arrays of descendant species exhibiting great ecological and phenotypic diversity (Schluter 2000). It is an important biotic feature of many oceanic archipelagos, and is especially prominent in Hawaiʻi, where conditions appear to favor unusually high rates of evolutionary innovation (Purruganan and Robichaux 2005, Givnish et al. 2009, Blonder et al. 2016). Invasion by alien species is a general problem affecting diversity worldwide, including on many oceanic archipelagos. Its impacts are especially severe in Hawaiʻi, where the native biota evolved in the absence of key guilds, such as large browsing mammals (Cuddihy and Stone 1990, Purugganan and Robichaux 2005).

The endemic Hawaiian silversword (Asteraceae) and lobeliad (Campanulaceae) lineages are two of the world’s premier examples of plant adaptive radiation (Robichaux et al. 1990, Baldwin 2003, Givnish et al. 2009, Blonder et al. 2016). The silversword lineage includes 33 species in three endemic genera (*Argyroxiphium, Dubautia*, and *Wilkesia*) (Carr 1985). The species grow in habitats as varied as exposed cinder and lava, dry shrubland, dry woodland, mesic forest, wet forest, and bog, and grow at elevations ranging from less than 100 m to more than 3,900 m (Robichaux et al. 1990). They also exhibit great phenotypic diversity, extending to most aspects of vegetative and reproductive form (Carr 1985, Purugganan and Robichaux 2005). They grow as rosette shrubs, small cushion-like shrubs, large shrubs, trees, and lianas, with leaves that range in length from as short as 5 mm to as long as 500 mm. Some species are monocarpic whereas others are polycarpic. Across species, the compound inflorescences range in size from small, with only three heads (capitulae), to massive, with up to 600 heads. Similarly, the heads contain as few as two flowers to as many as 650 flowers. Despite this exceptional ecological and phenotypic diversity, molecular phylogenetic analyses of nuclear rDNA ITS sequences provide compelling evidence that the silversword lineage is monophyletic (Baldwin and Robichaux 1995, Baldwin 2003), with a maximal age for the most recent common ancestor of only 5.2 ± 0.8 million years (Baldwin and Sanderson 1998), which closely approximates the age of Kauaʻi, the oldest current high island in the archipelago. Molecular evolutionary analyses of floral regulatory genes demonstrate that this common ancestor was allotetraploid, arising from ancient hybridization and whole genome duplication involving species within the “Madia” lineage of North American tarweeds (Barrier et al. 1999). The analyses also reveal that the rapid and extensive phenotypic diversification of the silversword lineage at the macroevolutionary scale has been accompanied by an accelerated rate of regulatory gene evolution (Barrier et al. 2001, Purruganan and Robichaux 2005). Molecular population genetic analyses further reveal that selection in contrasting ecological settings has been sufficiently strong to drive substantial quantitative divergence in key vegetative and reproductive traits at the microevolutionary scale in recently derived, parapatric species and to maintain the divergence in the face of significant gene flow detected at the level of individual regulatory genes (Lawton-Rauh et al. 2007) and more broadly across the genome (Remington and Robichaux 2007).

The lobeliad lineage includes 126 species in six genera (*Brighamia, Clermontia, Cyanea, Delissea, Lobelia*, and *Trematolobelia*) (Givnish et al. 2009). With its very high species richness, the lineage accounts for about one out of every eight species in the entire Hawaiian flora (Wagner et al. 1990, Givnish et al. 2009). Like the silversword lineage, the lobeliad lineage exhibits great ecological diversity, with the species growing in habitats as varied as exposed sea cliffs, dry woodland, mesic forest, wet forest, and bog (Givnish and Montgomery 2014, Scoffoni et al. 2015). The species also exhibit great phenotypic diversity, especially in growth habit, leaf size and form, and flower and fruit size and shape (Givnish et al. 2009, Givnish and Montgomery 2014). Molecular phylogenetic analyses provide strong evidence that the lobeliad lineage, like the silversword lineage, is monophyletic (Givnish et al. 2009).

Though the silversword and lobeliad lineages are marvels of evolutionary diversification, they and the Hawaiian flora more broadly are confronted by a suite of threats, especially from alien species. Browsing and trampling by alien ungulates pose severe threats to most native plants and ecosystems in Hawaiʻi (Cuddihy and Stone 1990), decimating even highly abundant, keystone elements of some landscapes (Robichaux et al. 1998). Key alien ungulates in Hawaiʻi include feral pigs (*Sus scrofa*) deriving from early Polynesian and European introductions, and feral goats (*Capra hircus*), sheep (*Ovis aries*), and cattle (*Bos taurus*) deriving from early European introductions (Cuddihy and Stone 1990, Nogueira-Filho et al. 2009). Additionally, mouflon sheep (*O. aries* subsp. *musimon*), axis deer (*Axis axis*), and mule deer (*Odocoileus hemionus*) have been introduced more recently as game animals on some islands (e.g., mouflon sheep on Hawaiʻi Island) (Cuddihy and Stone 1990). Competition by alien plants also poses severe threats, with some alien plants having the potential to dominate and transform native ecosystems (e.g., *Psidium cattleianum*), including dramatically altering fire regimes (e.g., alien grasses) and nutrient regimes (e.g., *Morella faya*) (Vitousek and Walker 1989, Loh and Daehler 2008, D’Antonio et al. 2011). Additional key threats to native plant reproduction include: (1) loss of native insect pollinators because of predation by alien social insects (e.g., Argentine ants (*Linepithema humile*) and yellowjacket wasps (*Vespula pensylvanica*)) (Wilson et al. 2009, Hartley et al. 2010), (2) loss of native bird pollinators and seed dispersers because of infection by alien *Plasmodium relictum* and *Avipoxvirus* spp., which are transmitted by alien mosquitoes (e.g., *Culex quinquefasciatus*) and cause avian malaria and pox, respectively (Pratt 2005, Atkinson and LaPointe 2009), and (3) predation of seeds and seedlings by alien rodents (e.g., black rats (*Rattus rattus*)) and slugs (e.g., *Limax maximus*) (Cuddihy and Stone 1990, Joe and Daehler 2008, Pender et al. 2013). (The Hawaiian biota includes no native social insects (Wilson 1996) and no native land mammals, except bats (Cuddihy and Stone 1990).)

Combined with other threats (e.g., habitat conversion associated with human settlement, agriculture, logging, and ranching) (Cuddihy and Stone 1990), these myriad alien threats have caused precipitous declines in the distributions and abundances of many Hawaiian plants. More than 50% of all Federally listed endangered flowering plants are endemic to Hawaiʻi (USFWS 2016), even though the archipelago accounts for less than 0.2% of the total U. S. land area. The silversword and lobeliad lineages exemplify the daunting magnitude of the problem. More than 20% of the species in the silversword lineage and more than 40% of the species in the lobeliad lineage are listed as endangered (USFWS 2016). Additionally, at least two species in the silversword lineage (Carr 1985, Wood 2015) and at least 20 species in the lobeliad lineage (Wagner et al. 1990, USFWS 2008a, Wood 2015) appear to have gone extinct.

In this paper, we highlight our sustained approach to addressing this daunting problem. We first discuss in detail our collaborative reintroduction efforts over the past 20 yr with the endangered Kaʻū silversword and Pele lobeliad in Hawaiʻi Volcanoes National Park (Fig. 1), efforts that have involved rediscovery, managed breeding, outplanting, and (for the Kaʻū silversword) facilitated achene dispersal following flowering. We next discuss linking the reintroduction efforts to landscape restoration at large scales in the Park and in adjacent State and private lands, thereby increasing the opportunities for substantial population growth and expansion of the Kaʻū silversword and Pele lobeliad in the future. We then discuss extending the reintroduction efforts, including the link to landscape restoration, to encompass all other endangered silversword and lobeliad taxa occurring historically on the eastern slopes of Mauna Loa or on Kīlauea. The ultimate objective of our sustained approach is to restore the possibility of adaptive radiation of the silversword and lobeliad lineages going forward, especially on the youngest and most geologically active, and thus perhaps most evolutionarily dynamic, part of the Hawaiian archipelago.

**Figure 1.**
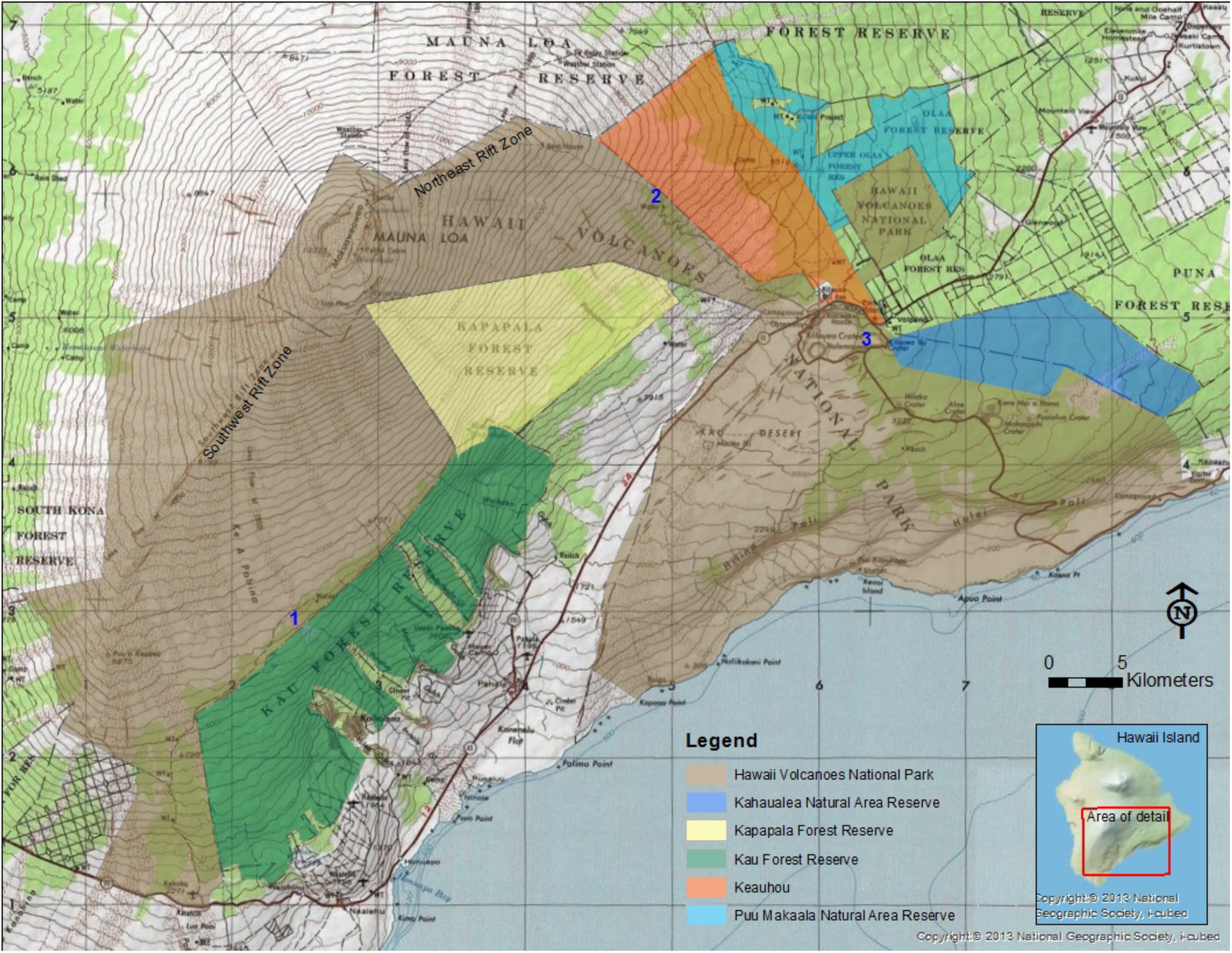
Portions of Mauna Loa and Kīlauea on Hawaiʻi Island, with Hawaiʻi Volcanoes National Park and adjacent State and private lands. Dark blue numbers denote areas in the Park in which we have reintroduced the Kaʻū silversword or Pele lobeliad. 1: Kahuku, 2: Kīpuka Kulalio, 3: Nāhuku. (Reintroduced populations of both taxa occur in the general vicinity of 1, but in different habitats on differently aged lava flows.)

## Reintroduction of the Endangered Kaʻū Silversword in Hawaiʻi Volcanoes National Park

*Argyroxiphium kauense* is a caulescent rosette shrub with long, narrow, and highly pubescent leaves (Carr 1985). Plants typically have a single rosette and are monocarpic. After growing for 15-20 or more years, the rosettes produce a large compound inflorescence with many heads. The rosettes of large individuals may exceed 70 cm in diameter and produce compound inflorescences exceeding 2 m in length and containing up to 300 heads, each with up to 300 flowers. Most flowering occurs from June to August. The plants are self incompatible (Carr 1985, USFWS 1996). The flowers are insect pollinated, primarily by yellow-faced bees (*Hylaeus* spp.). The achenes (single-seeded fruits) are mainly wind dispersed. At peak flowering, the plants are striking in appearance (Fig. 2), giving rise to their iconic status in Hawaiʻi.

**Figure 2.**
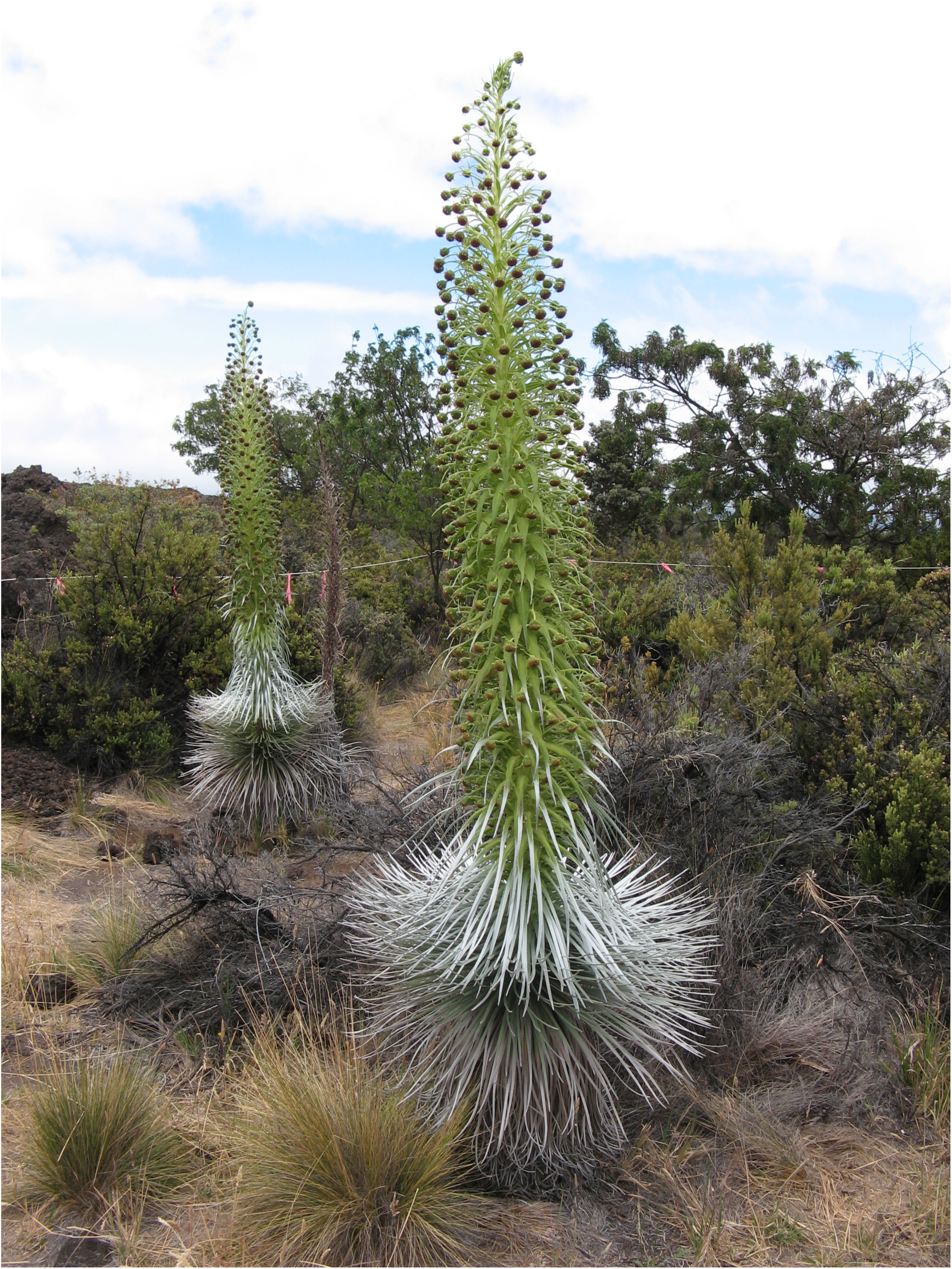
Flowering Kaʻū silverswords at the Kīpuka Kulalio site in Hawaiʻi Volcanoes National Park. The plants were outplanted as seedlings in 2000 and flowered in 2012. The large rosette of the plant in the foreground exceeds 70 cm in diameter and its massive compound inflorescence exceeds 2 m in length. The habitat is montane mesic-to-dry shrubland with scattered trees. The dead shrubs between the two flowering Kaʻū silverswords are a legacy of the severe drought of 2010, when total May-to-August rainfall recorded at the closest RAWS site was only 14.2 mm (see text). (All photos are by R. H. Robichaux.)

*Argyroxiphium kauense* exists in two distinctive morphological forms, with the forms having different ecological and geographical distributions. The Kaʻū form (or Kaʻū silversword) occurred historically in montane open mesic forest and mesic-to-dry shrubland habitats from about 1,600 to 2,600 m elevation in a broad band extending between the active southwest and northeast rift zones on Mauna Loa. At present, there are only two small remnant populations of the Kaʻū silversword. By contrast, the Waiākea form (or Waiākea silversword) occurred historically in montane bog and adjacent wet forest habitats from about 1,600 to 1,900 m elevation to the north of the northeast rift zone on Mauna Loa. At present, there is only one small remnant population of the Waiākea silversword. The significant divergence in form between the Kaʻū and Waiākea silverswords is evident when individuals are grown side-by-side in a common greenhouse environment (Fig. 3).

**Figure 3.**
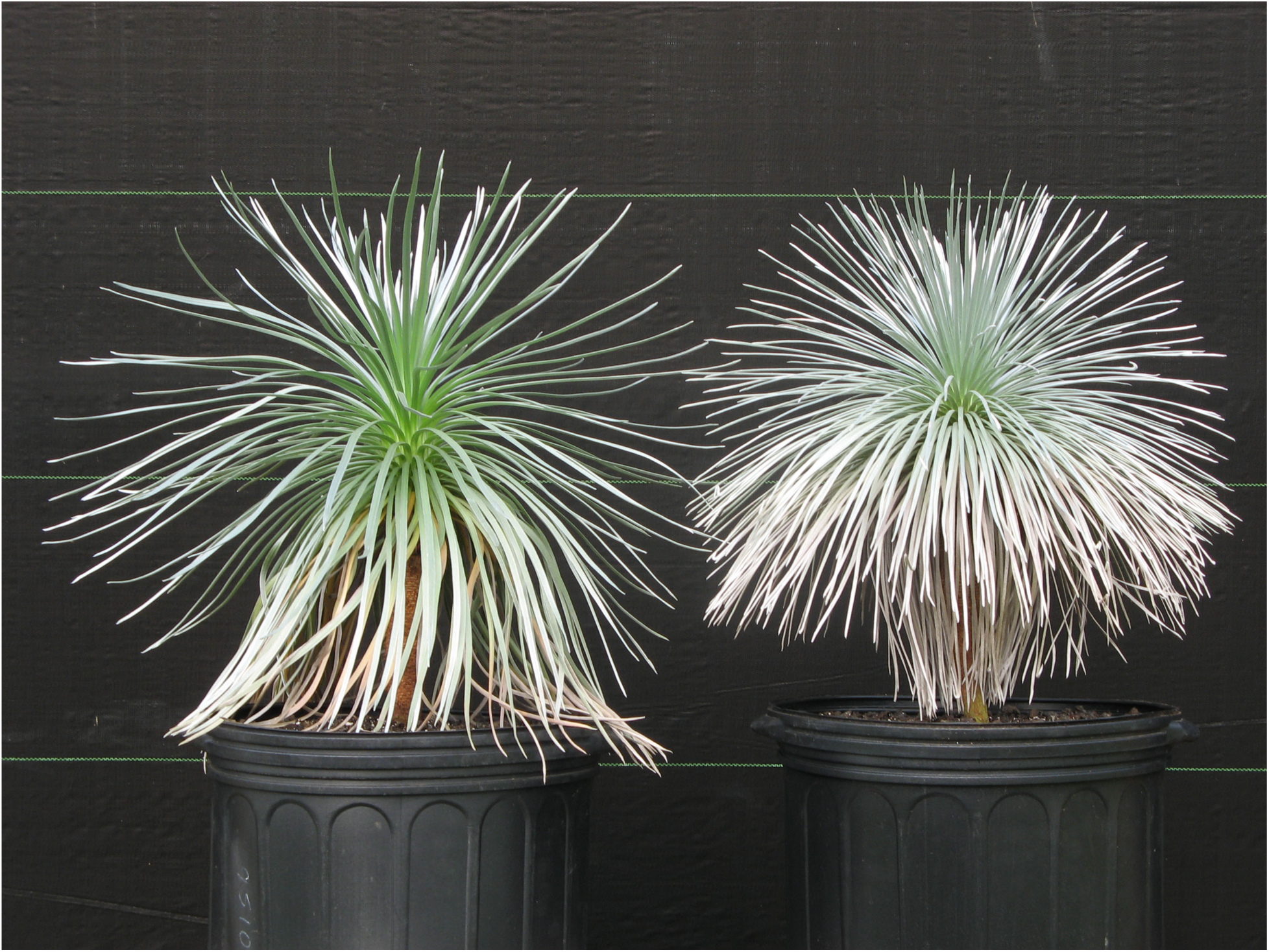
Kaʻū silversword (right) and Waiākea silversword (left) growing side-by-side at the Volcano Rare Plant Facility. For scale, the horizontal lines on the backdrop are separated by 30 cm.

Early descriptions, frequent sightings, and collections suggest that the Kaʻū silversword was widely distributed across the landscape prior to the impacts of alien ungulates, with at least some of the populations containing many thousands of individuals (USFWS 1996). Because of ungulate browsing and trampling, however, the Kaʻū silversword declined precipitously in distribution and abundance (USFWS 1996), such that only two small remnant populations persist within the historical range.

One remnant population occurs in the Kahuku region near the southern end of the historical range, in montane open mesic forest habitat. The area in which the population occurs, which includes the type locality for *A. kauense*, was privately owned and managed as Kahuku Ranch until 2003, when it was acquired by the Park. (The acquisition enlarged the Park by more than 61,000 ha.) A 1-2 ha exclosure was constructed in 1982 by the Kahuku Ranch foreman around part of the remnant population to provide some protection from alien ungulates (USFWS 1996). The exclosure was upgraded in the 1990s by the Hawaiʻi Division of Forestry and Wildlife working collaboratively with Kahuku Ranch (USFWS 1996). Following the Park’s acquisition of the area, the exclosure was again upgraded in 2004, using a 2-m tall fence design to more effectively exclude alien ungulates, especially mouflon sheep. A total of 381 remnant plants survive within the exclosure, with rosette diameters ranging from 4 to 70 cm.

The other remnant population occurs in the Kapāpala region in the central portion of the historical range, in montane mesic-to-dry shrubland habitat with scattered trees. By the late 1970s, the population was known to include only about 100 plants, all with small-to-medium rosette diameters (USFWS 1996). By the middle 1990s, the population was considered possibly to have been extirpated, especially with the high numbers of alien ungulates, including expanding mouflon sheep populations, in the region (USFWS 1996). However, in October 1996, we were re-exploring the Kapāpala region when we found two very small and previously browsed Kaʻū silverswords with aged main stems (Fig. 4). In 1997, 1998, and 1999, we revisited the area and exhaustively surveyed it. We discovered and retrieved 127 diminutive remnant plants, all about the size of the plant in Fig. 4 or smaller. We transported the plants by helicopter to the Volcano Rare Plant Facility. Because it was difficult to extract the root systems from the pāhoehoe lava substrate, many of the retrieved plants had few-to-no attached roots (i.e., they were essentially cuttings). Nonetheless, 96 plants established in pots, grew to large size, flowered, and produced viable achenes within 1-4 yr at the Rare Plant Facility. Additionally, the Park fencing crew, working collaboratively with the Hawaiʻi Division of Forestry and Wildlife, constructed a 4 ha exclosure (with a 2-m tall fence) in 1999 in the area, which is State land. Six remnant plants survive within the exclosure, with none having grown appreciably or flowered since 1999.

**Figure 4.**
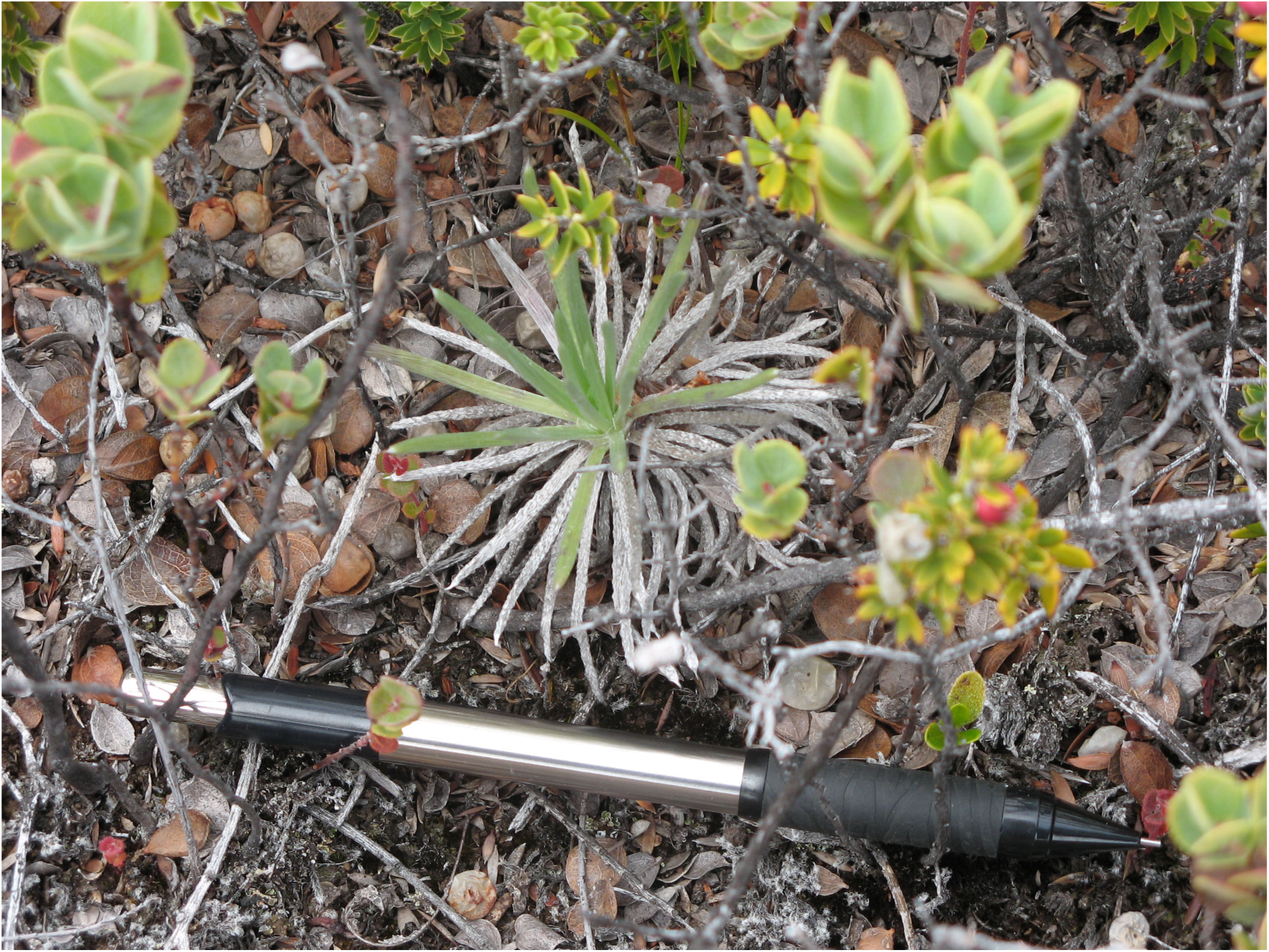
Remnant Kaʻū silversword rediscovered in the Kapāpala region in 1996, with a mechanical pencil for scale.

We have used the two remnant populations as source populations for reintroduction efforts in two areas of the Park on Mauna Loa. The remnant Kahuku population has served as the source for reintroduction in the Kahuku area of the Park, whereas the remnant Kapāpala population has served as the source for reintroduction in the Kīpuka Kulalio area of the Park (Fig. 1). Both areas of the Park have been the focus of landscape restoration at large scales. The Kahuku area is significantly wetter than the Kīpuka Kulalio area (Giambelluca et al. 2013), with the latter also having a more pronounced summer dry season. The reintroduction site in Kahuku consists of montane open mesic forest habitat, whereas the reintroduction site in Kīpuka Kulalio consists of montane mesic-to-dry shrubland habitat with scattered trees. Our reintroduction efforts at both sites have entailed two sequential phases, with the first involving managed breeding and outplanting, and the second involving facilitated achene dispersal.

### Reintroduction in Kahuku

We conducted managed breeding at the Rare Plant Facility and in the remnant Kahuku population. We established a managed breeding population at the Rare Plant Facility consisting mainly of plants that we retrieved from the remnant Kahuku population in 2004. Patterned after our successful retrieval of Kapāpala plants, we focused on a set of small but old plants in the remnant Kahuku population. As for the Kapāpala plants, many of the retrieved Kahuku plants were essentially cuttings because of the difficulty in extracting the root systems from the lava substrate. These retrieved plants were augmented in the managed breeding population at the Rare Plant Facility by several plants deriving from achenes that we collected from flowering remnant plants in the late 1990s, prior to the Park’s acquisition of the area. Under cultivation, the plants grew to large size and flowered. A total of 37 plants flowered at the Rare Plant Facility, mainly in 2006 and 2008. The remnant Kahuku population, unlike the remnant Kapāpala population, includes larger plants. A total of 36 larger remnant plants flowered in the population in 2004 and 2005, enabling us to expand our managed breeding efforts to the field.

For plants flowering at the Rare Plant Facility, we collected pollen in bulk from all other plants flowering at the same time, mixed it thoroughly, and applied it with adjustable cosmetic blush brushes to the stigmas on recipient heads. We used bulk pollen in order to increase the genetic diversity of offspring for each maternal founder. For plants flowering in the remnant population, we followed the same protocol, adjusting it as needed to overcome the challenges of rain and wind and to minimize the impacts of our pollen collecting on foraging by yellow-faced bees.

We collected, dried, and stored the mature achenes in a refrigerator at 4 °C. For each of the 73 maternal founders, we conducted a germination trial to assess percent viability of the achenes, then used the latter information to do targeted achene sowing at the Rare Plant Facility to produce seedlings for outplanting (Fig. 5). Our goal was to ensure that all maternal founders were represented in the reintroduced population, but that no maternal founder accounted for more than 2.5% of the total number of seedlings (i.e., that no maternal founder was substantially over-represented). Following the propagation protocol described in Moriyasu and Robichaux (2003), we grew the seedlings for 6-9 mo at the Rare Plant Facility, at which point they were typically 10-12 cm in height.

**Figure 5.**
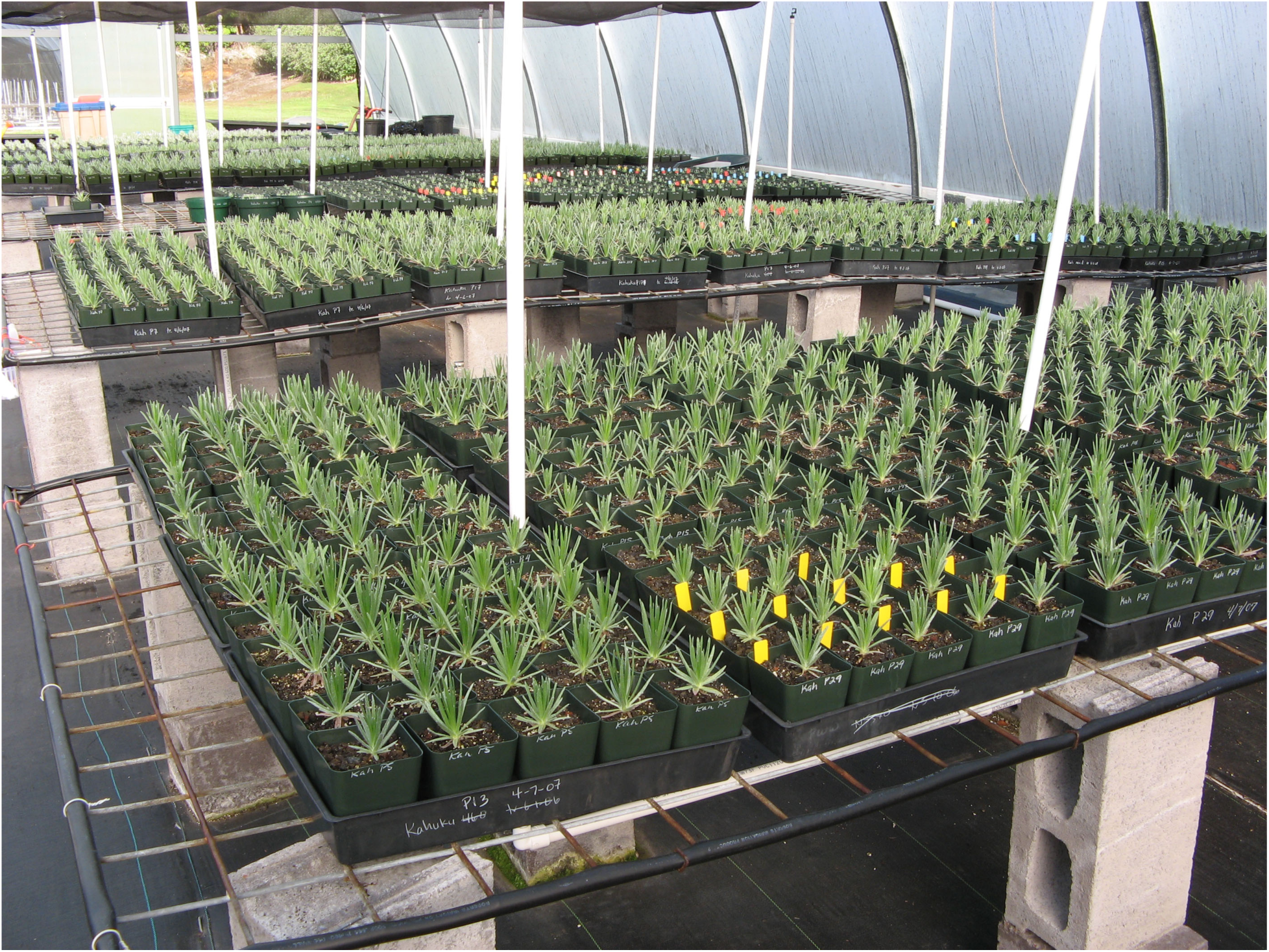
Kaʻū silversword seedlings at the Volcano Rare Plant Facility. The seedlings are destined for outplanting at the Kahuku site in Hawaiʻi Volcanoes National Park. Each seedling is tracked by maternal founder.

We outplanted the seedlings at a site in the Kahuku area of the Park centered at about 1,850 m elevation. The core part of the site, where we outplanted 95% of the seedlings, includes a 12 ha exclosure and 4 ha exclosure (with 2-m tall fences) and spans 1.3 km in cross-slope distance. The small remnant population occurs a short distance away from the core part of the site. Because of the highly patchy soil distribution on the lava substrate, especially with respect to soil depth, we used pin flags to identify individual planting locations for the seedlings prior to outplanting, with the planting locations preferably having a soil depth greater than 15 cm. We transported the seedlings to the remote site in large Rubbermaid ActionPackers, with each ActionPacker containing a mixture of seedlings from multiple maternal founders, which aided in their dispersion across the site. We watered the seedlings immediately after outplanting, then again about 2-4 wk later if rainfall during that initial period was low.

We outplanted 10,212 seedlings from the 73 maternal founders at the site between 2004 and 2009. The maximal number of seedlings outplanted per maternal founder was 245. We monitored survival and growth of the seedlings for the first decade from the start of outplanting (i.e., from 2004 to 2014), then shifted our focus to annual monitoring for flowering and new seedling recruitment. As of late summer 2014, 5,894 outplanted individuals were alive in the reintroduced population, with many of the plants having grown to more than 40 cm in rosette diameter (Fig. 6).

**Figure 6.**
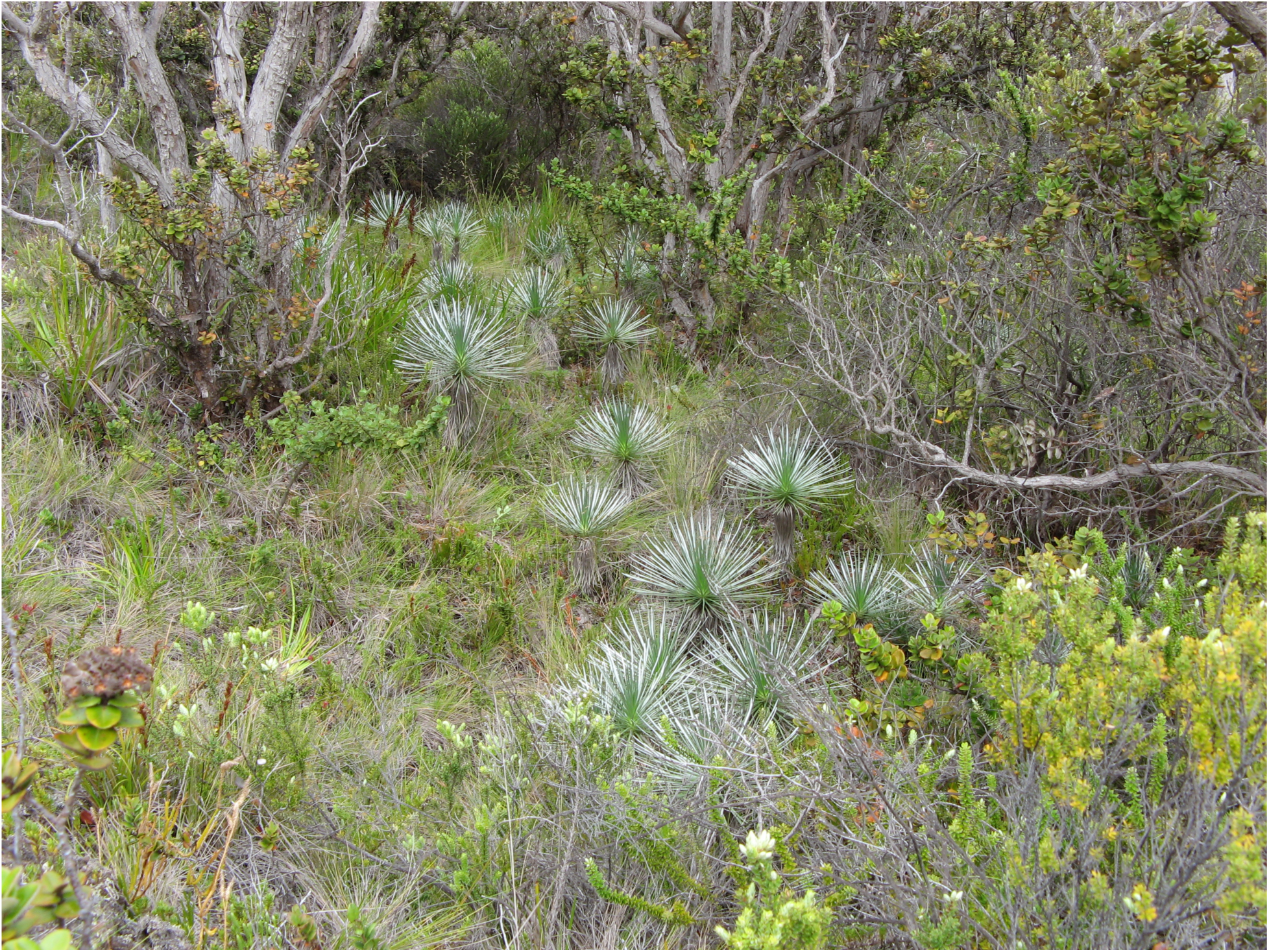
Kaʻū silverswords at the Kahuku site in Hawaiʻi Volcanoes National Park. The plants were outplanted as seedlings in 2007 and 2008. The rosettes of the larger plants in the photo approach 50 cm in diameter. The habitat is montane open mesic forest.

The first flowering in the reintroduced population was in 2014, when 46 plants flowered. An additional seven plants flowered in 2015 and 58 plants in 2016. We observed abundant visitation of the flowering plants by yellow-faced bees, often with 50 or more bees simultaneously visiting different heads on a single large compound inflorescence. Once the achenes were mature, we collected and dispersed them widely in the reintroduced population (i.e., as facilitated achene dispersal), while also allowing some to disperse naturally from the parent plants. Also as part of our facilitated achene dispersal effort to grow the reintroduced population, we collected and dispersed achenes widely from 57 remnant plants (or new founders) that flowered in the field after we had completed the outplanting phase of the reintroduction effort, with most of them flowering in 2014 and 2016. As of late summer 2016, 2,845 seedlings had established from the facilitated and natural achene dispersal in the reintroduced population.

### Reintroduction in Kīpuka Kulalio

We conducted managed breeding only at the Rare Plant Facility, with the managed breeding population consisting of plants that we retrieved from the remnant Kapāpala population in 1997 to 1999. A total of 96 plants flowered at the Rare Plant Facility, mainly in 1999 to 2001. Our protocols for hand pollinating the plants, collecting and sowing the achenes, and growing the seedlings for outplanting were the same as discussed above.

We outplanted the seedlings at a site in the Kīpuka Kulalio area of the Park centered at about 2,150 m elevation. The core part of the site, where we outplanted 96% of the seedlings, includes a 12 ha exclosure and 8 ha exclosure (with 2-m tall fences) and spans 1.3 km in cross-slope distance. We used pin flags to identify individual planting locations for the seedlings prior to outplanting and transported the seedlings to the site in ActionPackers. We watered the seedlings immediately after outplanting, then 1-3 additional times at 2-4 wk intervals if rainfall during that initial period was low.

We outplanted 11,060 seedlings from the 96 maternal founders at the site between 2000 and 2005. The maximal number of seedlings outplanted per maternal founder was 242. As in Kahuku, we monitored survival and growth of the seedlings for the first decade from the start of outplanting (i.e., from 2000 to 2010), then shifted our focus to annual monitoring for flowering and new seedling recruitment. As of late summer 2010, only 165 outplanted individuals were alive in the reintroduced population, although many of the surviving plants, especially those growing in flow channels in the lava with deeper soil pockets, had grown to large size, some with rosette diameters exceeding 70 cm (Fig. 2).

The first flowering in the reintroduced population was in 2007, when one plant flowered. A total of 69 additional plants flowered in subsequent years, primarily in 2012, 2014, 2015, and 2016. As in Kahuku, we observed abundant visitation of the flowering plants by yellow-faced bees. We collected and dispersed the mature achenes widely in the reintroduced population, while also allowing some to disperse naturally from the parent plants. As of late summer 2016, 578 seedlings had established from the facilitated and natural achene dispersal in the reintroduced population, with some seedlings having reached moderate size (Fig. 7).

**Figure 7.**
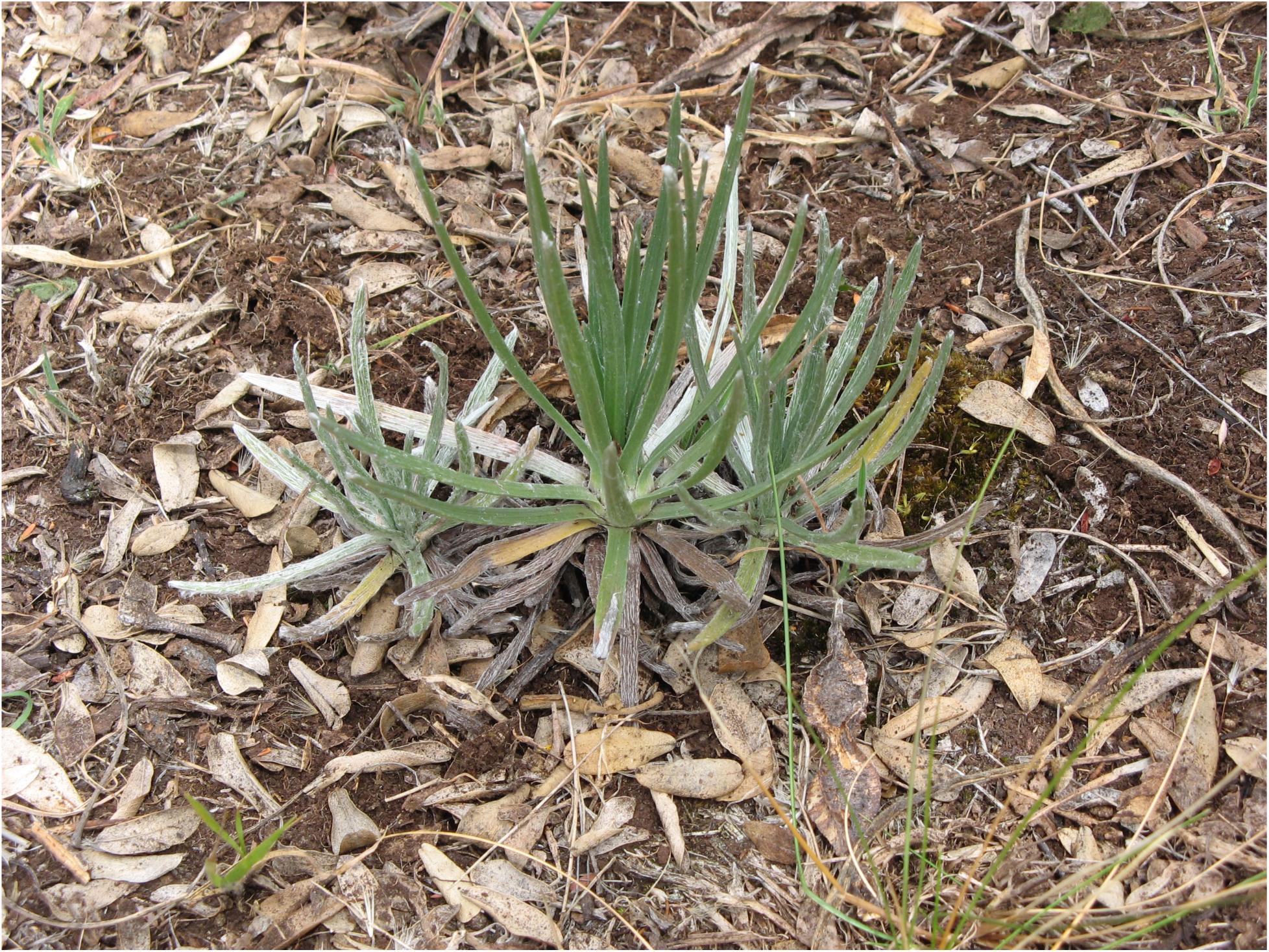
Kaʻū silversword seedling establishment following flowering at the Kīpuka Kulalio site in Hawaiʻi Volcanoes National Park. The largest of the three adjacent seedlings is 7-8 cm in height.

The recruitment of new seedlings in the reintroduced population coincides with an increase in summer rainfall in the Kīpuka Kulalio area of the Park in recent years. Data from the closest Interagency Remote Automatic Weather Station (RAWS), which is located in the Park a short distance downslope from the core reintroduction site, indicate that total May-to-August rainfall each year between 2000 and 2013 ranged from 14.2 mm to 270.0 mm (mean = 166.1 mm) (NWCG 2016). By contrast, total May-to-August rainfall each year between 2014 and 2016 ranged from 531.9 mm to 577.6 mm (mean = 550.2 mm) (NWCG 2016), with multiple major tropical storms passing near or over the island each summer.

## Reintroduction of the Endangered Pele Lobeliad in Hawaiʻi Volcanoes National Park

*Clermontia peleana* is a large shrub to small tree of montane wet forest habitat (Rock 1913). Large individuals may reach 4-6 m in height. In addition to growing in soil, plants are able to grow epiphytically, especially on old-growth *Metrosideros polymorpha*, the dominant canopy tree. The tubular, decurved flowers produce copious nectar (Fig. 8) and are bird pollinated, mainly by curve-billed honeycreepers, several species of which (e.g., the Hawaiʻi mamo (*Drepanis pacifica*)) are now extinct (Pratt 2005). Most flowering occurs from May to September. In addition to outcrossing, the plants are capable of selfing. The very small seeds, which are contained in large orange fruits, are bird dispersed.

**Figure 8.**
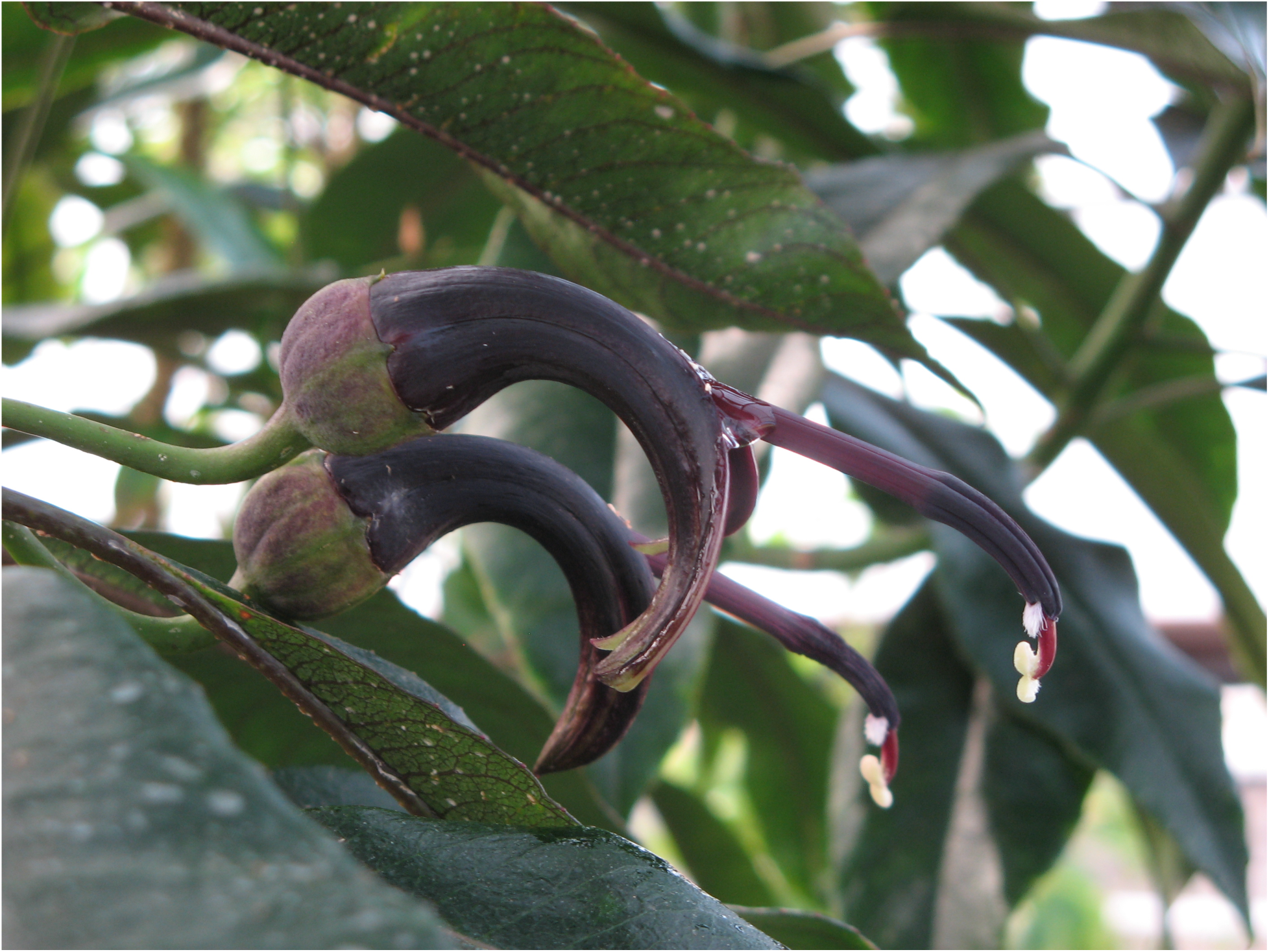
Flowering Pele lobeliad in the managed breeding population at the Volcano Rare Plant Facility. The tubular, decurved corollas are 8-10 cm in length. The flowers are protandrous. Initial elongation of the style within the staminal column causes pollen to accumulate near the tip of the column, immediately adjacent to the small tuft of white hairs visible on the flower in the foreground. When visiting honeycreepers (or perhaps other birds) brush against the tuft of hairs, a pore opens in the staminal column allowing abundant pollen to be released onto the heads and necks of the visitors. Further elongation of the style over 1-2 d causes the stigma to extend beyond the staminal column. The stigma then reflexes to expose its receptive surface, which is well-positioned to receive pollen from additional visiting honeycreepers. The flowers produce copious nectar, visible as a droplet at the junction of the corolla and staminal column in the flower in the foreground.

The predominant form of *C. peleana* is the blackish-purple flowered form. It is the form originally described by Rock (1913) and is the form to which we refer as the Pele lobeliad. It occurred historically on Kīlauea, the eastern slopes of Mauna Loa, and parts of Mauna Kea, with the type locality being about 6-8 km below the summit caldera of Kīlauea. A much less common form, differing only in having greenish-white rather than blackish-purple flowers, is represented in a very small number of the early collections (Lammers 1990).

By 2000, the last known individuals of the Pele lobeliad growing in the wild had died (Liittschwager and Middleton 2001, USFWS 2003, 2008b). At that time, the only known surviving individual was growing in cultivation at the Rare Plant Facility, where we had succeeded in germinating one seedling from an older seed collection and growing it to moderate size. However, in August 2007, we were re-exploring an area along Wailuku stream near the boundary between Mauna Loa and Mauna Kea substrates and discovered two remnant plants (Fig. 9). With multiple subsequent explorations of this and other areas, we found six remnant plants in total, all growing epiphytically in heavily degraded montane wet forest habitat with major alien ungulate damage and alien plant invasion in the understory.

**Figure 9.**
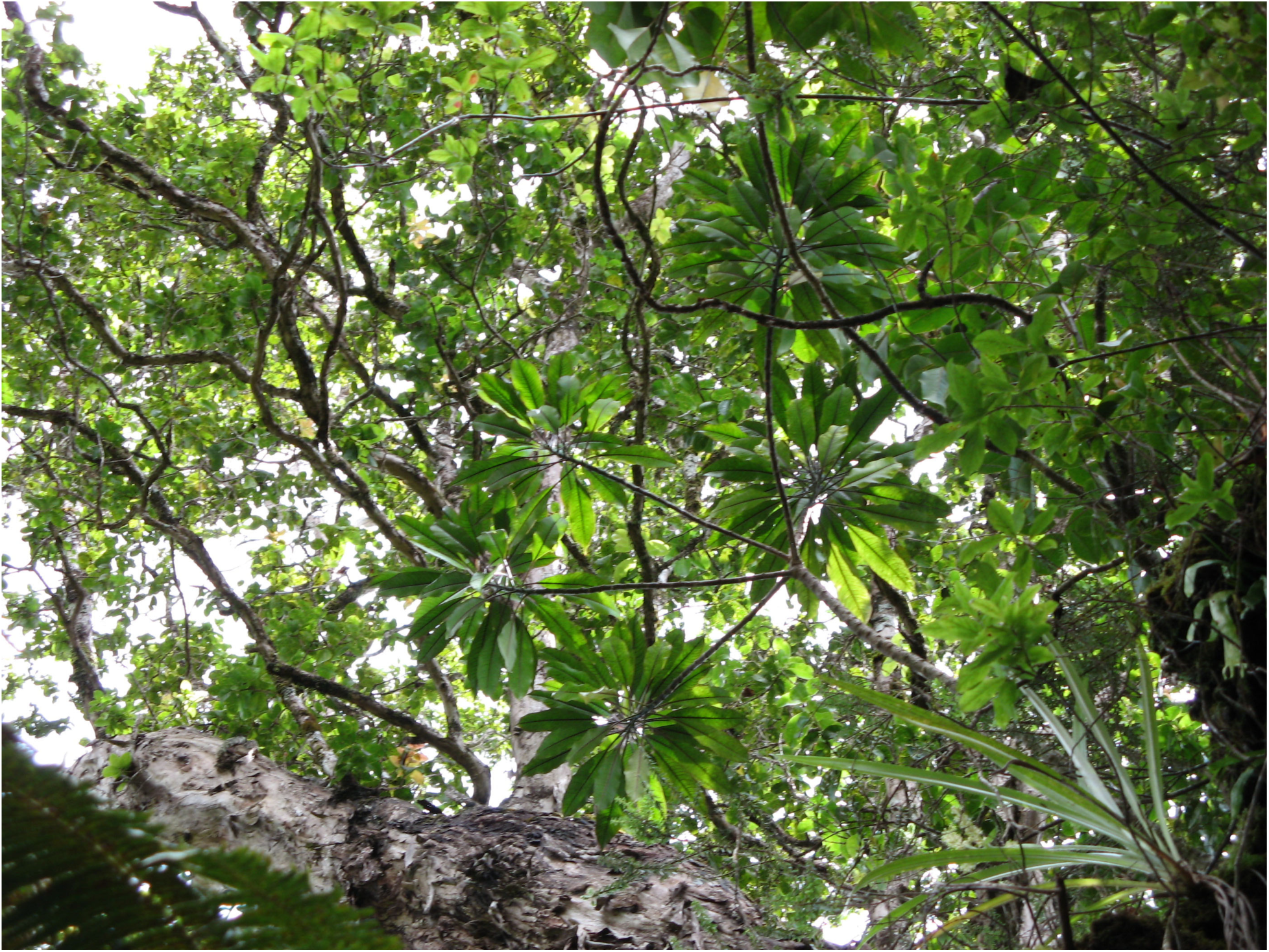
Remnant Pele lobeliad rediscovered in 2007. The plant (with the five shoot branches and large leaves) is growing epiphytically about 6 m up in the canopy of an old-growth *Metrosideros polymorpha*.

We conducted managed breeding of the Pele lobeliad, working initially with the first founder at the Rare Plant Facility. Our discovery of the six additional remnant plants increased the number of potential founders available to us, although only two of the plants were reproductively mature when we discovered them. Unfortunately, the six remnant plants grow at elevations sufficiently low to have alien mosquitoes that serve as vectors for avian malaria and pox (Pratt 2005, Atkinson and LaPointe 2009). Thus, few if any honeycreepers are present, which greatly reduces open pollination. Hand pollinating the remnant plants in the field proved to be difficult because of the abundant rains in the montane wet forest habitat, coupled with the fact that the plants grow epiphytically high in the forest canopy.

To help overcome these challenges, we adapted an air-layering technique used with tropical fruit trees in Hawaiʻi. For each air-layer, we carefully cut a section of the shoot a short distance below the terminal foliage, excising the bark down through the phloem (to block sugar transport) along 10-12 cm of shoot length. We wrapped the exposed portion of the shoot with moist sphagnum, then sealed it with Bemis Parafilm M film to protect the air-layer from the abundant rains. Following root proliferation at the terminal end of the air-layer, typically within 3-4 mo, we harvested the shoot (Fig. 10). The air-layering technique was challenging because we had to be suspended on climbing ropes high in the forest canopy while carefully cutting into the shoots with razors. Nonetheless, we successfully produced air-layers in 2010 on five of the six remnant plants, which then enabled us to grow them as rooted cuttings at the Rare Plant Facility. Each of the founders flowered in one or more subsequent years at the Rare Plant Facility, even those that were not yet reproductively mature in the field, which enabled us to hand pollinate them.

**Figure 10.**
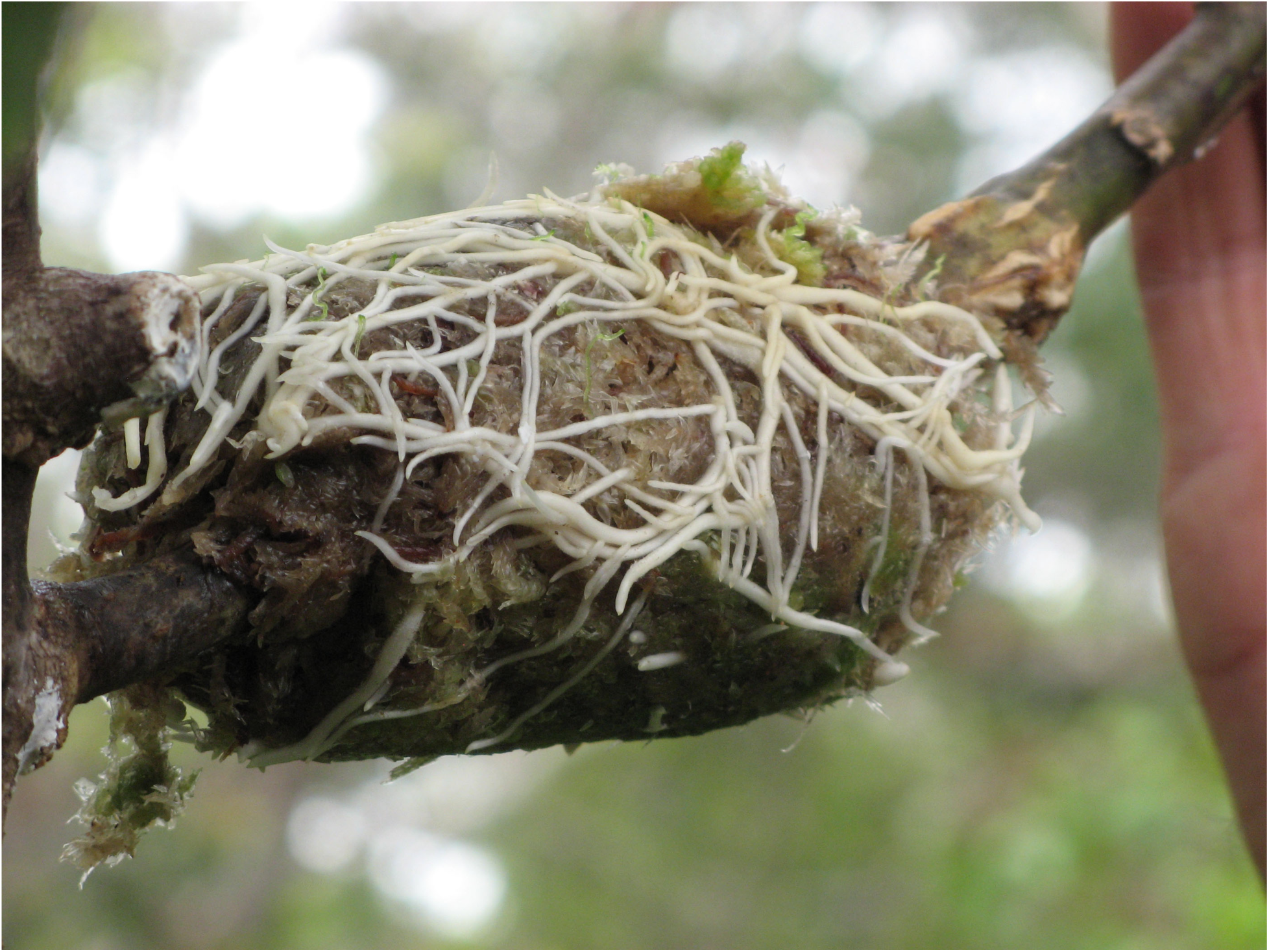
Roots from air-layering of a remnant Pele lobeliad. The dense mat of material surrounding the newly severed shoot is sphagnum used in the air-layering. The rooted shoot, which derives from the plant in Fig. 9, has subsequently flowered in the managed breeding population at the Volcano Rare Plant Facility.

We used very small brushes to transfer pollen between flowers. Because *Clermontia* flowers are protandrous (Fig. 8) (Aslan et al. 2014), we collected newly mature pollen from a flower ready to release it from the staminal column, then transferred the pollen to a different flower whose stigma had very recently emerged from the staminal column. Most such hand pollinated flowers produced mature fruits with some viable seeds. By contrast, most flowers that were not hand pollinated aborted without producing fruits. Though the number of flowers per founder was low, we generated multiple mature fruits via either selfing or outcrossing from each of the six founders growing at the Rare Plant Facility. We sowed the seeds immediately after harvesting and cleaning the fruits. We grew the seedlings following a propagation protocol similar to that for the Kaʻū silversword. We grew them for about 24 mo, at which point they were 40-45 cm in height.

Using the small set of known founders as our sole source population, we have reintroduced the Pele lobeliad in two areas of the Park on Mauna Loa and Kīlauea (Fig. 1). Both areas of the Park have been the focus of landscape restoration at large scales. The reintroduction sites in both Kahuku and Nāhuku consist of montane wet forest habitat. The Nāhuku site (at about 1,200 m elevation) is closer elevationally and geographically to the type locality for the Pele lobeliad (Rock 1913). It has a higher density of large tree ferns (*Cibotium* spp.) and thus a more closed canopy structure than the Kahuku site. The Kahuku site (at about 1,800 m elevation) is above the zone with alien mosquitoes, and thus has more abundant populations of honeycreepers and other native birds than the Nāhuku site (Gorresen et al. 2007). (Modeling of avian malaria and pox transmission along an altitudinal gradient on the eastern slopes of Mauna Loa and on Kīlauea has shown that disease transmission essentially disappears at elevations above about 1,500 m, where alien mosquito populations decline to very low levels, resulting in disease free, higher elevation “refugia” for honeycreepers and other native birds (van Riper et al. 1986, 2002, Atkinson and LaPointe 2009)).

### Reintroduction in Kahuku

The Kahuku site includes a 10 ha exclosure (with a 2-m tall fence), within which we established a grid of 36 evenly spaced 40-m diameter circular plots. We transported the seedlings to the remote site in ActionPackers. For each maternal founder, we outplanted the seedlings in an even distribution across the grid of plots. Soil moisture levels were typically high at the time of outplanting, such that supplemental watering was typically unnecessary to aid seedling establishment.

We outplanted 697 seedlings from five maternal founders at the site between 2013 and 2016, with the number of seedlings per maternal founder ranging from 80 to 225. As of late summer 2016, 613 outplanted individuals were alive in the reintroduced population, with some plants exceeding 2 m in height. The first flowering at the site was in 2016, when one plant flowered.

### Reintroduction in Nāhuku

The Nāhuku site has two separate grids of 4 and 8 evenly spaced 40-m diameter circular plots. The site spans 0.7 km in cross-slope distance. As at the Kahuku site, soil moisture levels were typically high at the time of outplanting, which precluded the need for supplemental watering.

We outplanted 110 seedlings from the first maternal founder at the site between 2009 and 2011. In 2015, we outplanted 203 seedlings from the five additional maternal founders, with the number of seedlings per maternal founder ranging from 14 to 65. As of spring 2016, 277 outplanted individuals were alive in the reintroduced population, with some plants exceeding 3 m in height. Unlike at the Kahuku site, no plants have yet flowered at the Nāhuku site.

## Reintroduction and its Link to Landscape Restoration on Mauna Loa and Kīlauea

We continue to pursue reintroduction efforts with the Kaʻū silversword and Pele lobeliad in the Park. The current focus of our efforts with the Kaʻū silversword is additional facilitated achene dispersal following flowering at the Kahuku and Kīpuka Kulalio sites. The current focus of our efforts with the Pele lobeliad is additional outplanting at the Kahuku and Nāhuku sites to provide for more balanced representation among the six maternal founders used to date, including adding seedlings from the first maternal founder at the Kahuku site, where that founder is not yet represented. We also expect to outplant seedlings at both sites within the next 3-5 yr from the seventh (and last known) maternal founder. We succeeded in 2012 in obtaining one viable seed from the one fruit produced in the field during the past nine years by the seventh maternal founder, and are now growing the resulting juvenile plant at the Rare Plant Facility. The latter founder is the most spatially isolated of the remnant plants. For reasons that are unclear, we have been unsuccessful in our air-layering efforts with it in the field. However, we are now using air-layering of the juvenile plant at the Rare Plant Facility to generate 2-3 additional individuals, and expect that one or more of them will flower within the next 1-3 yr, which should enable us to obtain seedlings for outplanting.

For both the Kaʻū silversword and Pele lobeliad, we have linked our reintroduction efforts to landscape restoration at large scales in the Park. As part of its multifaceted landscape restoration efforts (HAVO 2013, 2016), the Park has fenced large areas on Mauna Loa and Kīlauea, pioneering new construction protocols for ungulate-proof fences across challenging volcanic landscapes, including across expansive ʻaʻā and pāhoehoe lava flows with highly irregular surface features. The Park has eliminated alien ungulates within the fenced landscapes using a combination of ground-based and aerial-assisted methods (HAVO 2013). It also has implemented regular inspection and maintenance protocols for all fences to preclude new ungulate ingress. (Park fences on Mauna Loa and Kīlauea now total more than 256 km in length.) The Park has controlled alien plants, especially those with the potential to dominate and transform native ecosystems, in key areas using integrated mechanical and chemical methods. It also has implemented regular monitoring protocols to detect and remove new seedlings emerging either from alien seed banks or from bird dispersed alien seeds (e.g., of *Morella faya*) (Woodward et al. 1990, Loh and Daehler 2008) and new shoots emerging from persistent alien rhizomes (e.g., of *Hedychium gardnerianum*) (Minden et al. 2010). Additionally, the Park has implemented monitoring and sanitation protocols, including for field vehicles, equipment, and gear, to help prevent the introduction or spread of alien plants, social insects, and other organisms. Coupled with planting efforts involving key canopy and understory species in critical areas, the Park has made major strides toward restoring very large tracts of the landscape on the eastern slopes of Mauna Loa between the southwest and northeast rift zones and on Kīlauea, including near the summit caldera.

Of direct relevance to the Kaʻū silversword, the Park’s landscape restoration efforts have been sufficiently successful that the recently acquired large Kahuku area now has dramatically reduced numbers of alien ungulates and few alien plants in the upper elevations, and alien ungulate numbers should reach zero over the next few years. Likewise, the large Kīpuka Kulalio area now has no alien ungulates and few alien plants. Further, the Park’s monitoring and sanitation protocols have helped to prevent the introduction of alien ants into both areas. The lack of alien ants, especially predatory Argentine ants, likely contributes to the healthy pollinator populations, as the solitary, ground nesting yellow-faced bees are especially vulnerable to Argentine ant predation (Cole et al. 1992, Hartley et al. 2010). Thus, both large areas of the Park offer the potential for substantial population growth and expansion of the Kaʻū silversword in the future.

Of direct relevance to the Pele lobeliad, the Park’s landscape restoration efforts have been sufficiently successful that the recently acquired large Kahuku area, as noted above, now has dramatically reduced numbers of alien ungulates and few alien plants in the upper elevations. Likewise, the large Nāhuku area now has no alien ungulates. Alien plants, including some with the potential to transform the native montane wet forest habitat (e.g., *Morella faya* and *Hedychium gardnerianum*), occur in the Nāhuku area, but the Park’s sustained control efforts have reduced them to very low numbers. Thus, both large areas of the Park offer the potential for substantial population growth and expansion of the Pele lobeliad in the future. This is especially true for the Kahuku area, with its more abundant populations of honeycreepers and other native birds (i.e., nectarivores and frugivores) (Gorresen et al. 2007). The degree to which the more limited abundance of honeycreepers and other native birds in the lower elevation Nāhuku area will constrain population growth and expansion of the Pele lobeliad is unknown. In this regard, it also is unknown whether alien nectarivorous and frugivorous birds in the Nāhuku area might serve as partial replacements, as has been documented for smaller-flowered *Clermontia* species in nearby areas with montane wet forest habitat (Aslan et al. 2014).

Landscape restoration efforts also have been implemented in State and private lands adjacent to the Park under the umbrella of the Three Mountain Alliance, which is a Federal/State/private watershed partnership (TMA 2007). These landscape restoration efforts, illustrated by the following two examples, mean that future population expansion of both the Kaʻū silversword and Pele lobeliad may extend beyond the boundaries of the Park. First, the Keauhou area, which is private land owned by Kamehameha Schools, shares a significant boundary with the Park (Fig. 1). More than 11,700 ha in the Keauhou area have been fenced by the Park fencing crew and others working collaboratively with Kamehameha Schools. This includes the area immediately adjacent to the Kīpuka Kulalio site where we have reintroduced the Kaʻū silversword. Almost all alien ungulates have now been eliminated from the fenced landscape and alien plants are being controlled. Key parts of the landscape contain montane mesic-to-dry shrubland habitat suitable for the Kaʻū silversword, thus providing potential opportunities for population expansion. Second, Kaʻū Forest Reserve, which is State land, also shares a significant boundary with the Park (Fig. 1). An 800 ha fenced exclosure in upper Kaʻū Forest Reserve has recently been constructed by the Park fencing crew working collaboratively with the Hawaiʻi Division of Forestry and Wildlife. The 800 ha exclosure is the initial phase of a planned 4,800 ha fenced area in upper Kaʻū Forest Reserve (DLNR 2012). The Kahuku site where we have reintroduced the Pele lobeliad is immediately adjacent to the new exclosure. Alien ungulates are being eliminated within the exclosure, which has few alien plants at present. The exclosure contains extensive montane wet forest habitat with significant populations of honeycreepers and other native birds (Gorresen et al. 2007, DLNR 2012), and thus provides potential opportunities for population expansion of the Pele lobeliad.

### Implications for adaptive radiation of the silversword and lobeliad lineages

Landscape restoration at large scales in the Park and in adjacent State and private lands on Mauna Loa and Kīlauea is critical to the long-term success of our reintroduction efforts not only for the Kaʻū silversword and Pele lobeliad but also for other endangered silverswords and lobeliads. For example, the Rare Plant Facility has grown seedlings of all other endangered silversword and lobeliad taxa occurring historically on the eastern slopes of Mauna Loa or on Kīlauea, which has enabled us to begin implementing reintroduction efforts for all of them. These collaborative reintroduction efforts are patterned closely on our efforts with the Kaʻū silversword and Pele lobeliad. They are linked to landscape restoration not only in the areas discussed above but also in the large Puʻu Makaʻala and Kahaualeʻa Natural Area Reserves, which are State lands that share significant boundaries with the Park (Fig. 1) (TMA 2007, DLNR 2013). The respective silversword and lobeliad taxa (and the numbers of seedlings outplanted to date) include: the Waiākea silversword (5,673), *Clermontia lindseyana* (2,605), *Cyanea platyphylla* (72), *Cyanea shipmanii* (1,443), *Cyanea stictophylla* (466), and *Cyanea tritomantha* (216). (Seedlings of the lobeliad taxa have been grown at both the Rare Plant Facility and the Park’s greenhouse facility.)

Importantly over the longer term, this link to landscape restoration on the eastern slopes of Mauna Loa and on Kīlauea may allow for the wide range of ecological, genetic, and other processes underlying lineage diversification to operate at microevolutionary and macroevolutionary scales in the silversword and lobeliad lineages (Purugganan and Robichaux 2005, Blonder et al. 2016). We fully recognize that even with this link, alien species will continue to pose daunting challenges. Nonetheless, this link should help substantially in restoring the possibility of adaptive radiation of the silversword and lobeliad lineages going forward, especially on the youngest and most geologically active, and thus perhaps most evolutionarily dynamic, part of the Hawaiian archipelago.

## Acknowledgments

Our silversword and lobeliad reintroduction efforts have been funded primarily by the U. S. Fish and Wildlife Service and National Park Service. Our landscape restoration efforts have been funded by the National Park Service, U. S. Fish and Wildlife Service, Hawaiʻi Department of Land and Natural Resources, and Kamehameha Schools. We thank Hawaiʻi Volcanoes National Park staff (including Howard Hoshide, Allen Ramos, Jon Faford, Keola Medeiros, Susan Dale, Caitlin French, and all members of the crews responsible for fencing, alien ungulate control, alien plant control, and native plant propagation), Hawaiʻi Division of Forestry and Wildlife and Natural Area Reserves System staff, Kamehameha Schools staff, Three Mountain Alliance staff, and David Okita of Volcano Helicopters for their expert assistance with our reintroduction and restoration efforts. We also thank Junior Emicilio of Kahuku Ranch and Nelson Santos of the Hawaiʻi Division of Forestry and Wildlife for their early fencing of the remnant Kahuku population of the Kaʻū silversword, Marta Leppes, Hoʻala Fraiola, and Steven Perlman for their assistance with our initial exploring for remnant individuals of the Kaʻū silversword and Pele lobeliad, Lance Tominaga for his assistance with technical climbing in the forest canopy and air-layering of the Pele lobeliad, Nicole Kuamoʻo for her early assistance with growing Kaʻū silversword seedlings at the Volcano Rare Plant Facility, Karin Schlappa for her assistance in drafting Figure 1 and providing summary data on fencing in the Keauhou area, and the large number of agency and community volunteers for their assistance with outplanting silversword and lobeliad seedlings.

## Author Contributions

We have implemented all of the efforts described in this paper collaboratively, with extensive sharing of expertise and effort. Our complementary lead responsibilities for the different phases are: (i) *rediscovering the remnant populations*: Robichaux and Warshauer for the Kapāpala population of the Kaʻū silversword; Bio, Robichaux, and Perry for the Pele lobeliad; (ii) *surveying the remnant populations and retrieving small plants*: Robichaux, Moriyasu, Bakutis, Perry, and Bergfeld for the Kahuku population and Robichaux, Warshauer, Perry, Bergfeld, and Tunison for the Kapāpala population of the Kaʻū silversword; (iii) *air-layering*: Enoka and Bio, with guidance from Moriyasu, for the Pele lobeliad; (iv) *growing adult plants for managed breeding and seedlings for outplanting at the Rare Plant Facility*: Moriyasu, Enoka, and Robichaux for the Kaʻū silversword and Pele lobeliad; (v) *managed breeding*: Robichaux and Moriyasu for the Kaʻū silversword; Moriyasu, Enoka, and Robichaux for the Pele lobeliad; (vi) *coordinating outplanting and monitoring:* Robichaux and Bakutis for the Kaʻū silversword; Robichaux, McDaniel, and Wasser for the Pele lobeliad; (vii) *facilitated dispersing of achenes*: Robichaux and McDaniel for the Kaʻū silversword; (viii) *coordinating landscape restoration:* Loh and Tunison in the Park; C. Cole, Rubenstein, and Robichaux collaboratively with Whitehead in the Keauhou area; Bergfeld and C. Cole in Kaʻū Forest Reserve; Agorastos and I. Cole in Puʻu Makaʻala and Kahaualeʻa Natural Area Reserves; and (ix) *extending the reintroduction efforts to other endangered silversword and lobeliad taxa*: Moriyasu and Enoka for growing seedlings at the Rare Plant Facility; all authors for collecting seeds from remnant plants or for outplanting.

